# Temperature and predation alter metabolic scaling without changing size-based structure community in freshwater macroinvertebrates

**DOI:** 10.1101/2025.06.27.661898

**Authors:** Vojsava Gjoni, Douglas S. Glazier, Justin Pomeranz, James Junker, Aria Smith, Jacob Woelber, Staci Reynolds, Trevor Welch, Jeff S. Wesner

## Abstract

Body size is a key trait that influences ecological processes such as metabolism, abundance, and species interactions. While the metabolic theory of ecology (MTE) proposes a universal scaling of metabolic rate with body mass, recent evidence shows that this relationship is not fixed. Environmental factors like temperature and predation can alter the metabolic scaling exponent, potentially reshaping size distribution. However, most research has examined these patterns within individual species, leaving open questions about how environmental drivers affect scaling at the community level. To address this, we performed a mesocosm experiment manipulating both temperature and fish predator presence in freshwater macroinvertebrate communities. We found that metabolic scaling at the community level is highly responsive to environmental context: warming steepened the scaling exponent in predator-free tanks but flattened it when predators were present. This suggests that larger individuals reduce their baseline metabolic rates under predation risk, especially at higher temperatures. Interestingly, the slope of the community size distribution remained stable across treatments, indicating that shifts in metabolic scaling occurred independently of changes in size structure. Together, these findings highlight the environmental sensitivity of metabolic scaling and suggest that links between metabolism scaling and size distribution may be more complex than MTE predicts.

## 1. Introduction

Body size plays a pivotal role in shaping ecological dynamics, influencing key processes such as metabolism, reproduction, and species interactions across biological scales (1–3). A central framework explaining these patterns is the metabolic theory of ecology (MTE), which posits that an organism’s metabolic rate (*R*) scales with body mass (*M*) following a power-law relationship: *R* = *aM^b^*. The exponent *b* is generally expected to approximate ¾, a value theoretically derived from models describing internal resource transport networks (4). This scaling rule has been widely used to predict how energy flows through individuals, populations and entire ecosystems (1,5).

Nonetheless, empirical work has increasingly shown that the exponent *b* is far from universal. It often varies across taxa, environments, and life stages (6–10). In aquatic macroinvertebrates, for instance, metabolic scaling can shift in response to both temperature and predation pressure, often becoming shallower than the canonical ¾ expectation (9,11,12). These observations suggest that ecological conditions can significantly influence metabolic allometry, revealing flexibility in a trait once considered physically constrained.

Variability in metabolic scaling has potentially important implications for the Individual Size Distribution (ISD) —the well-established pattern where smaller organisms tend to occur at higher densities than larger ones (13,14). Assuming energy use is equal across size classes, shifts in *b* should map directly onto changes in size spectra under equilibrium conditions (7,15,16). The ISD is described by a power law, *N ∼ M ^λ^*, where *N* is abundance and *M* is individual body mass. Metabolic scaling theory predicts that the exponent *λ* is generated by interactions of three community-level variables: trophic transfer efficiency (TTR), predator-prey mass ratio (PPMR), and metabolic scaling (*b*) (17–20), such that:

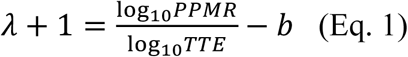

Given typical values of *PPRM* as 10^4^, of *TTE* as 0.1 and of *b* as 0.75, the exponent *λ* tends to be negative, with values typically ∼-1.75 to -2, underscoring a remarkably consistent ecological pattern across Earth’s diverse ecosystems (21,22). Yet, the link between the value of the metabolic scaling exponent *b* and the ISD’s *λ* has been rarely studied experimentally. Recent work demonstrated that even within a single species, evolutionary changes in body size can decouple metabolic scaling from abundance patterns, underscoring how energetics and demography interact in more nuanced ways than previously assumed (16).

Environmental temperature adds further complexity to this picture. Rising temperatures not only elevate metabolic demand through basic biochemical processes (23), at least up to a point before decreasing (24), but also they can shift the body-mass scaling relationship itself via size-specific physiological and behavioral responses (8,9,12). Moreover, temperature influences body size more directly through well-documented plastic responses, such as the temperature–size rule (25–27). Warmer environments are predicted to favor smaller organisms (28) leading to more negative values of *λ* (i.e., “steeper” slopes) and this has been observed empirically (29–32). Consequently, these size reductions may influence λ, linking thermal conditions to both metabolic scaling and ISD (16).

Predation, especially when it targets larger individuals, adds an important layer of ecological pressure. In many systems, larger prey face a greater risk of being eaten, often driving populations and communities toward smaller average body sizes (33,34). These changes go beyond simple shifts in size distributions; they also influence the way metabolic rates scale with body mass, by favoring traits that lower energy demands, such as reduced activity and slower growth in larger individuals. While the metabolic rates we measured reflect resting conditions, they are tightly connected to an individual’s overall capacity for movement, growth, and other energy-intensive functions, and are therefore sensitive to ecological pressures. Studies of amphipods have shown that exposure to higher temperatures and predator presence triggers size-specific adjustments: larger individuals tend to suppress energetically costly behaviors to avoid detection, while smaller, less visible individuals remain more metabolically responsive (9,12). This suggests a clear adaptive pattern. In environments with active predators, large individuals often restrict activity and growth— especially at higher temperatures when encounters are more frequent (35)—leading to a dampened metabolic response among adults. Meanwhile, smaller individuals, under less threat, continue to ramp up metabolic activity with warming, producing a shallower metabolic scaling (*b*). In predator-free settings, this pattern flips: small individuals appear to suppress metabolic responses, possibly due to risk of cannibalism from larger conspecific individuals, while adults take advantage of safer conditions to increase their metabolic rates. This reversal causes *b* to rise with temperature, a trend particularly evident in amphipods (9,12,36).

Although the effects of temperature and predation on metabolic scaling have been explored extensively, these investigations have largely focused on single species or isolated populations. What remains unclear is how these two drivers interact to influence metabolic scaling at the community level, where species vary in body size, life history, and ecological roles. Crucially, there is still a lack of experimental studies that directly examine how changes in metabolic scaling correspond to shifts in ISD under combined thermal and predation pressures. This missing link limits our understanding of how physiological processes scale up to influence community structure (16,37). As a result, we still lack empirical evidence about whether and how environmentally driven changes in metabolism propagate to reshape ISD at the community scale. Addressing this gap is essential to understand how temperature and predation jointly structure energy flow and body size patterns in natural systems.

To address this gap, we conducted a mesocosm experiment manipulating temperature and predator presence in freshwater macroinvertebrate communities. Our aim was to evaluate how these environmental drivers affect community-level metabolic scaling exponents and in turn, community body-size distributions. Specifically, we tested: (i) how does ecological context influence the metabolic scaling exponent *b* at the community scale? and (ii) do changes in *b* predict shifts in the ISD’s λ? By linking individual-level metabolism, which underlies community-level metabolic scaling, with body size distributions that shape size spectra, our work provides a valuable test of how physiological and structural patterns may emerge at the community scale. This approach offers a more ecologically grounded perspective on the scaling of metabolism and population abundance under environmental change.

## 2. Materials and methods

### Study Site

The experiment was conducted at the University of South Dakota’s Experimental Aquatic Research Site, Vermillion, SD. The site consists of 24 fiberglass tanks (1136 L). The tanks were filled with 714 L of water over a mix of cobble and sand for substrate in May of 2023. Water levels were monitored and maintained over the course of the experiment. Each tank had an overflow spout and a magnetic drive water pump (Danner Supreme Aqua-Mag). to ensure mixing and to bubble the surface to maintain sufficient O2. After filling, tanks were inoculated with ∼ 100ml of filtered (250 µm) local river water to introduce plankton.

Populations of insects colonized the tanks via oviposition beginning ∼1 day after they were filled and continuing through the experiment. AQQA Aquarium 800W Heaters (AQQA Inc.) were placed in half of the tanks and set to ∼5 degrees C above ambient nighttime temperatures. Temperature data were taken daily in the middle of the water column using a YSI Pro-Series DSS (YSI Inc., Yellow Springs, OH, USA). Heaters were adjusted to maintain a temperature of ∼5 degrees C above ambient (Figure S1). When averaged across dates, the heated mesocosms were warmer than the unheated mesocosms by 4.3 ± 1.9°C. Fish (*Lepomis cyanellus*) were gathered from a local pond with a seine net and set free in half of the tanks on 2023-05-10, six days after filling the tanks (one fish per tank). At the beginning of the experiment, their lengths ranged from roughly 8 to 10 cm, and by the end, they had grown to between 10 and 12.5 cm. The setup included four treatment combinations: ambient temperature with fish, ambient temperature without fish, heated tanks with fish, and heated tanks without fish (Figure 1).

**Figure 1.**
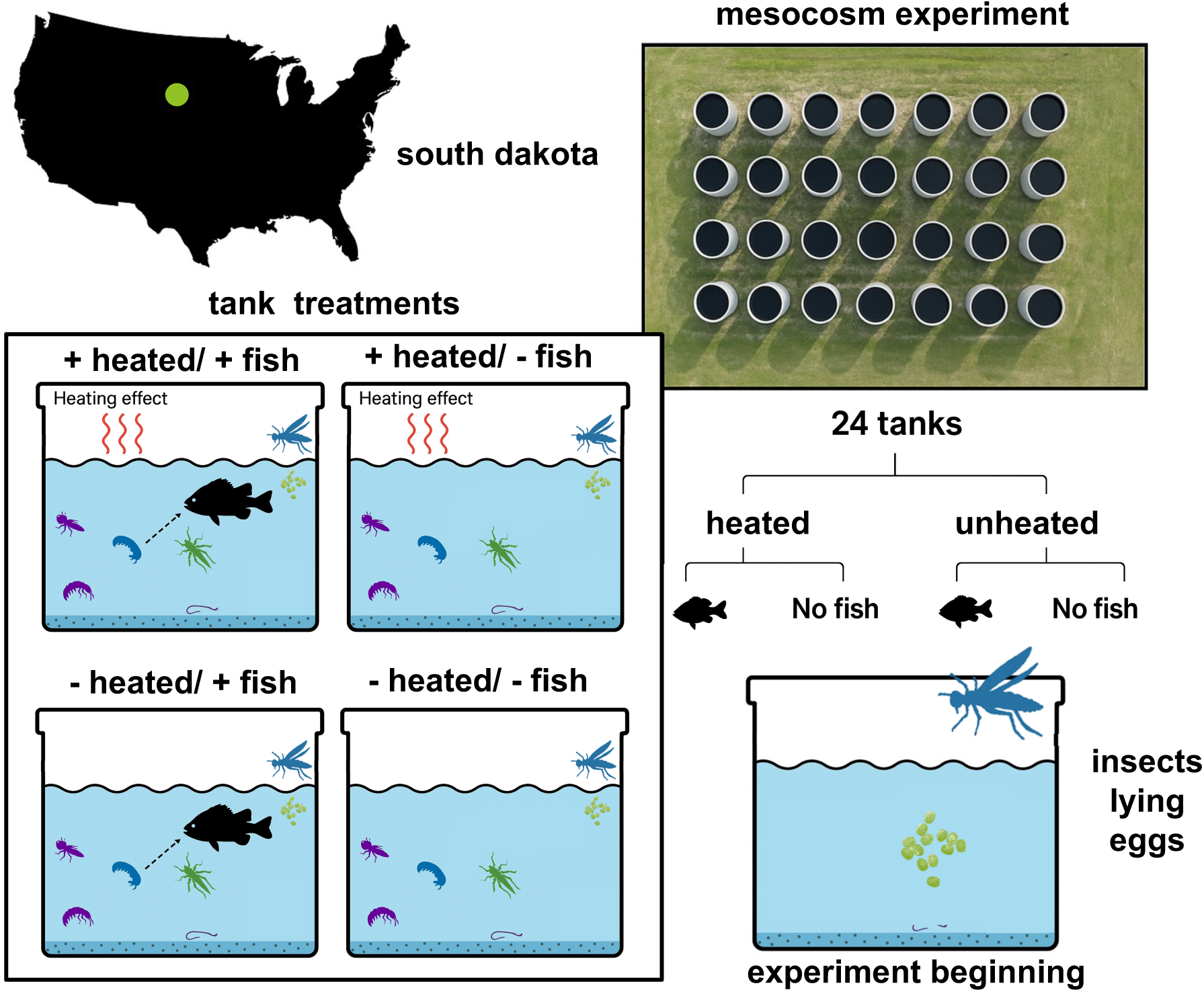
Overview of the experimental design. The study was conducted in South Dakota, USA (green dot on map), using 24 outdoor field mesocosms arranged in a 4 × 6 grid (aerial image, top right). Tanks were assigned to a fully factorial design crossing two treatments: temperature (heated vs. unheated) and fish presence (with vs. without *Lepomis cyanellus*), resulting in four treatment combinations. Schematics depict each treatment combination, with red wavy arrows indicating heated conditions and silhouettes of *L. cyanellus* representing fish presence. Macroinvertebrate prey of varying sizes and taxa naturally colonized the tanks. At the start of the experiment, aquatic insects oviposited into the tanks (bottom right), initiating community assembly. This design allowed us to test the interactive effects of temperature and predation on community-level metabolic scaling and size spectra.

### Macroinvertebrates

The 24 mesocosm tanks supported naturally assembling freshwater invertebrate communities. These communities developed from colonization by adult aquatic insects that used the tanks to lay their eggs, resulting in the emergence of a diverse and self-assembling assemblage. On average, each tank contained around seven taxa, with most taxa consistently present across all tanks. The communities included a range of freshwater invertebrates such as midges, beetles, and various odonate larvae (e.g., damselflies and dragonflies), several of which are known aquatic predators.

To investigate patterns in ISD, we measured a total of 1,529 individuals across these mesocosms. Tanks with fewer than 100 individuals were excluded from the ISD analysis to ensure sufficient sample size and data quality, resulting in the exclusion of three tanks. For the metabolic scaling analysis, 415 individuals were measured, with approximately 50 individuals sampled from each of three replicate tanks per treatment. Sampling was designed to include all species present within each tank and to capture a representative range of body sizes. This ensured robust comparisons of metabolic scaling across treatments, accounting for the diversity and size structure of the invertebrate communities.

### Metabolic rate measurements

Metabolic measurements occurred 30 days after treatments were started. A Hess sampler (0.032 m^2^, 500 um collection mesh) was used to sample benthic macroinvertebrates following the methods of (26). Macroinvertebrates from the sediment were sorted and placed in containers with filtered water (Whatman GF/F 0.7 *μ*m glass fibre filters to remove metabolically active microbes) from their corresponding treatment and held without food for 24 hours to empty their guts. After 24 hours, they were separated into individual glass vials with 20 mL of oxygenated, filtered water. Nine experimental vials with a macroinvertebrate and one control without (to correct for oxygen changes not due to respiration) were used for each run of measurements. Two O2 readings were taken an hour apart with each vial using a Fibox 4 oxygen sensor (Presens, Regensburg, Germany). The macroinvertebrates were then photographed (lengths measured using ImageJ) before being dried and measured on a microbalance. Oxygen consumption was estimated using the following equation:

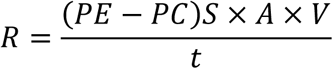

*R* is the oxygen consumption*, PE* and *PC* represent partial pressures of oxygen in the experimental environment (20 mL vial with an invertebrate) and control, respectively, *S* is the solubility coefficient of oxygen in water, *A* is the volume of 1 mol O2 at standard temperature and pressure, *V* is the volume of water in the vials, and t is the time of incubation.

To ensure accurate results, preliminary testing was performed to see how long it would take for a visible drop in dissolved oxygen content to occur. This determined the 60-minute time span between measurements. During the experiment, macroinvertebrates were kept in water that corresponded to their respective thermal treatment of predation and temperature.

### Individual Size Distribution

On the last day of the experiment (after 30 days), we collected benthic macroinvertebrates with a 0.09 m^2^ Hess sampler. The sampler was forced through the sediment and rock substrate so it was flush with the bottom of the mesocosm. Then we disturbed the substrate by hand for 10 seconds and flushed the substrate through a 250 𝜇m mesh net (38,39). The sample was then placed on a 250 𝜇m sieve, large debris (e.g., rocks) was scrubbed and removed, and the remaining sample was preserved in 95% EtOH. To measure body sizes of benthic macroinvertebrates, we sorted a subsample and retained the first 200-500 individuals. This approximates the number of individuals in simulation studies for which estimates of individual size distributions stabilize (39). These individuals were placed on a gridded watch glass, photographed, and measured for length to the nearest pixel using ImageJ. The pixels were converted to mm using the grid as a reference and adjusting for changes in magnification. Then the dry mass (mg) of each individual was determined using family-specific length-mass equations (Benke et al. 1999).

### Data Analysis

To estimate the community-level metabolic scaling exponent *b*, we fit log-transformed respiration as a function of log-transformed body mass using a Gaussian linear mixed model. Each individual’s respiration rate was measured and matched to its body mass, and these individual-level data were nested within mesocosm tanks, which served as the unit of replication. The model included a three-way interaction among log body mass, temperature, and predator treatment, with tank identity as a varying intercept. This structure allowed us to estimate tank-level scaling relationships while accounting for differences in intercepts across replicates. The prior for the scaling exponent was set to Normal(0.75, 0.2) to reflect prior assumptions of the ¾ scaling pattern. The remaining regression coefficients were set to Normal(0,1) and the intercept was Normal(0,0.2). Both log respiration and log mass were mean-centered prior to analysis to facilitate model convergence and align priors with the centered data. The resulting community-level value of *b* thus reflects the average slope across tanks, while variation in intercepts captures differences in baseline metabolic rates across communities.

To estimate ISD’ *λ*, we fit a generalized linear mixed model with individual body size as the response variable, heat and fish treatment and their interactions as the predictor variables, and tank as the varying intercept. The likelihood was a truncated Pareto, which allows the model to estimate coefficients reflecting the frequency distribution of the individual body sizes *λ* (40). Additional information required for the truncated Pareto are xmin and xmax, which represent the minimum and maximum body sizes from each tank. To set xmin, we used the *estimate_xmin()* function from the *poweRlaw* package, which determines the minimum body size for which the data still follow a power law. For xmax we used the measured maximum body size in each tank. Importantly, fish as organisms were excluded from all analyses of size distribution; only invertebrate body sizes were used to estimate *λ*.

We fit the models using *brms* (41) and *isdbayes* (40) in R version 4.4.2 (R Core Team 2024). Posteriors were explored with NUTS sampling in Stan (Stan Development Team 2024). We used 2000 iterations and 4 chains for each model. Model convergence was checked via plotting the chains and ensuring that R-hat was less than 1.01. Model fit was checked using posterior predictive checking (42) (Figure S2, Figure S3). Prior implications were checked using prior predictive checking (43) (Figure S4).

### Simulation Study

To determine how likely our empirical estimates of *λ* and *b* met the assumption of the MTE we used equation 1 to simulate *λ* across a range of plausible values of *PPMR* (0.05 to 0.4), *TTE* (10^2^ to 10^7^) and *b* (0.3, 0.75, 1) and compared those values to empirical *λ*s from the mesocosms.

### Data Availability

All data and code are publicly available at https://github.com/jswesner/NEON-2023-mesocosm-experiment (to be permanently archived on Zenodo upon acceptance).

## 2. Results

Our results support the hypothesis that warming increases the metabolic scaling exponent when predator risk is absent, but that this effect is reversed under predation pressure. However, the lack of change in *λ* across treatments suggests that shifts in individual-level energetics may not directly translate into changes in community-level size structure, indicating a possible disconnect between metabolic scaling and emergent ISD patterns.

### Metabolic scaling

Metabolic scaling exponents were responsive to both heating and predation treatments (Figure 2a; Table 1). In the presence of fish predators, heating caused a clear reduction in the metabolic scaling exponent, from 0.46 under ambient conditions to 0.35 under heated conditions (Table 1), indicating an >99% probability of decline. This pattern is consistent with a suppression of metabolic rate in larger individuals, as previously hypothesized in predator-sensitive taxa. In contrast, when fish were absent, heating increased the metabolic scaling exponent substantially, from 0.31 at ambient temperature to 0.45 under heated conditions (Table 1), indicating a >99% probability of an increase. These opposing trends demonstrate a strong interactive effect of temperature and predation on the metabolic scaling relationship.

**Figure 2.**
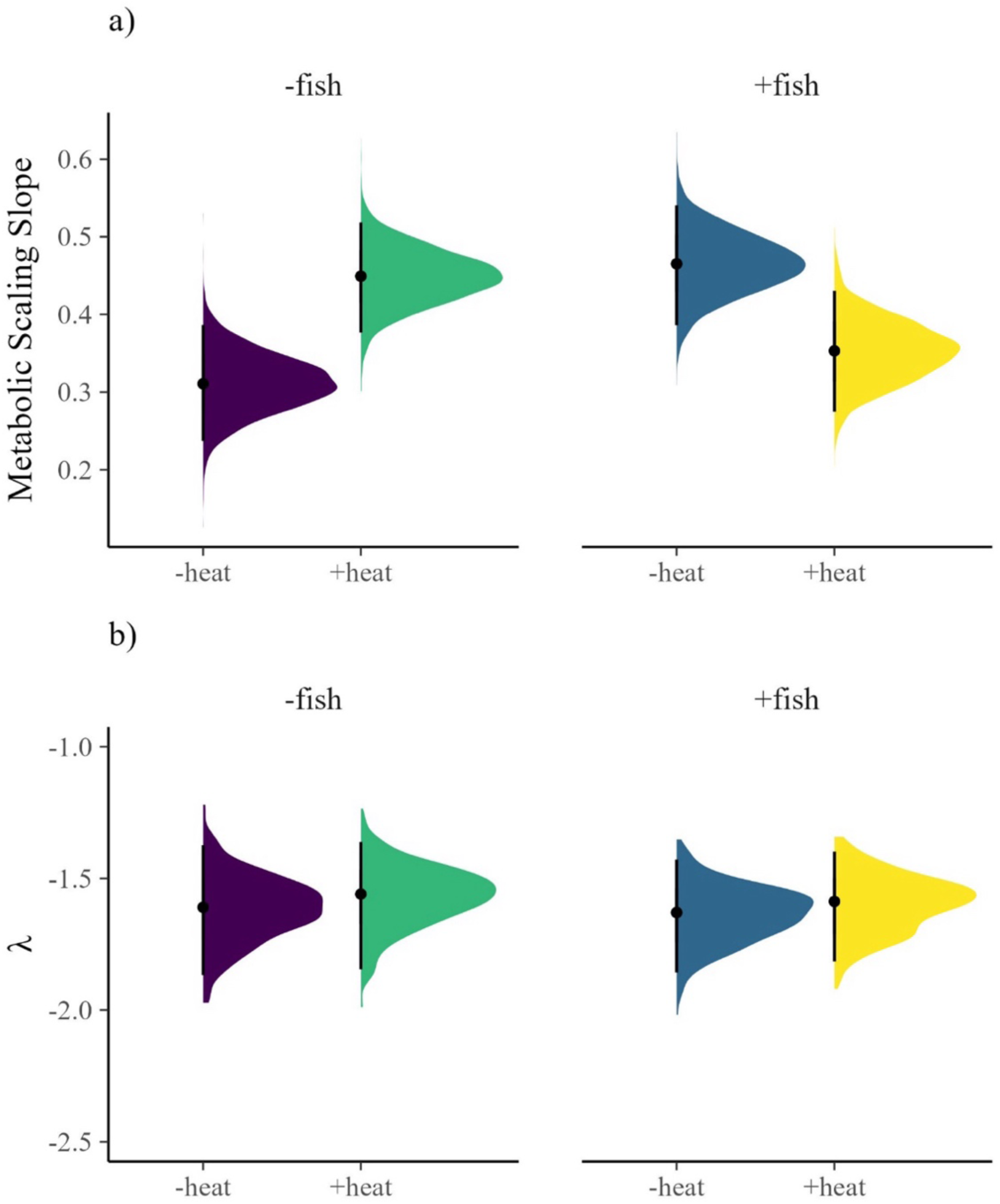
Horizontal density plots show the full posterior distributions of (a) the metabolic scaling slope and (b) the ISD (*λ*) across four treatments in a factorial experiment (Ambient vs. Heated temperature, crossed with presence vs. absence of fish predators). Points represent posterior means, and error bars show 95% credible intervals. Color indicates treatment group. While heating did not significantly affect *λ* (panel b), it did alter the metabolic scaling *b* (panel a), with contrasting effects depending on predation regime. Specifically, heating reduced the metabolic scaling slope in the presence of fish but increased it in their absence, indicating interactive effects of temperature and predation on metabolic allometry, independent of changes in community size structure.

**Table 1.**
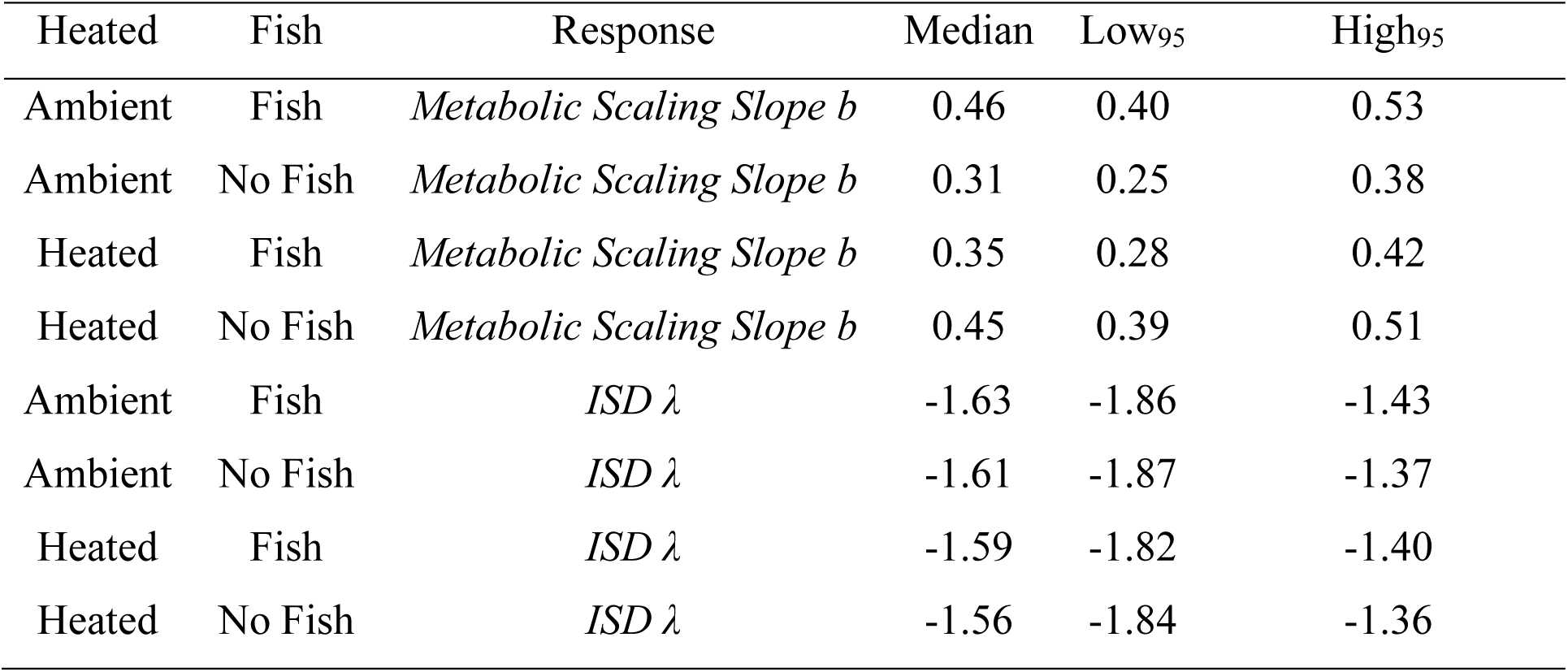
Median and 95% credible intervals for the metabolic scaling slope *b* and the size-abundance parameter λ estimated from the individual size distribution (ISD).

### Individual size distribution

The slope of the ISD’s 𝜆 remained largely consistent across both temperature and predation treatments (Figure 2b; Table 1). Across all experimental conditions, 𝜆 values ranged narrowly between –1.63 and –1.56, with overlapping credible intervals indicating no meaningful differences (Table 1). Specifically, neither warming nor the presence of fish predators altered the value of 𝜆. This suggests that, despite changes in community temperature and predator cues, the overall structure of abundance relative to body size remained stable. Thus, while temperature and predation clearly influenced individual metabolic scaling (see below), these effects did not directly translate into detectable shifts in community-level size–abundance patterns.

### Simulation Study

Equation 1 indicates that *λ* is shaped by the interplay among trophic transfer efficiency (*TTE*), predator–prey mass ratio (*PPMR*), and the metabolic scaling exponent *b*. However, variation in these parameters does not necessarily lead to shifts in *λ*. Simulations from equation 1 across a wide range of *TTE* and *PPMR* values show that our empirical *λ*s are most plausible when *b* is <0.75 (Figure 3). Notably, empirical *λ* values from our study overlapped closely with the simulation outputs where *b* ranged between 0.3 and 0.5 and *TTE* remained moderate. This suggests that the observed invariance in *λ* across environmental treatments could be explained by compensatory shifts in food web parameters that buffer the effects of metabolic scaling changes.

**Figure 3.**
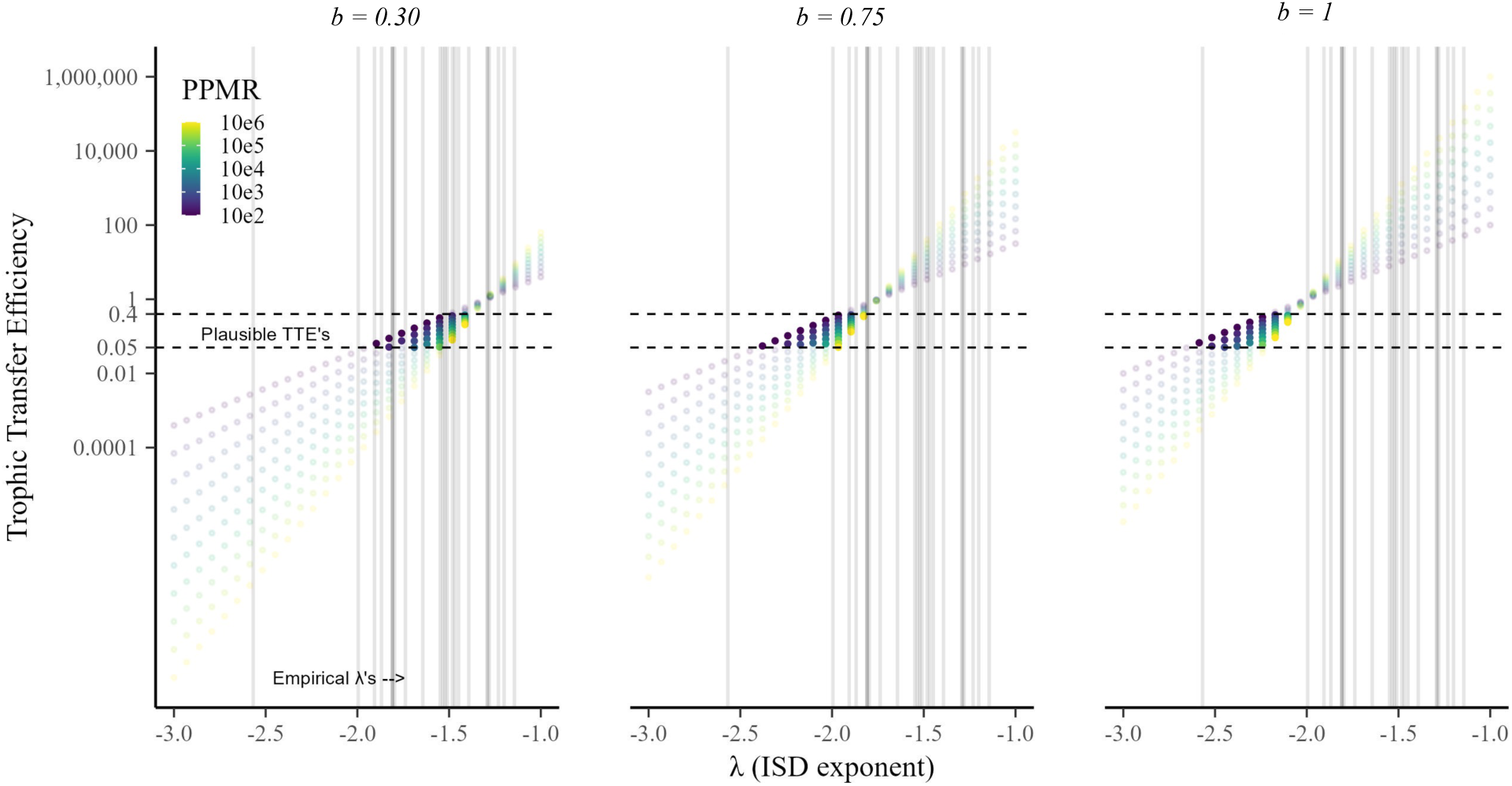
Simulated effects of metabolic scaling exponent *b*, trophic transfer efficiency (*TTE*), and predator–prey mass ratio (*PPMR*) on the ISD slope (*λ*). Panels show combinations of *λ* values across different *b* values (0.3, 0.75, and 1). Each colored point represents a unique pairing of *TTE* and *PPMR*, with color indicating *PPMR* magnitude. Horizontal dashed lines mark the range of ecologically plausible *TTE* values, while vertical gray lines indicate empirically observed *λ* values. The figure demonstrates that multiple parameter combinations can yield similar *λ* outcomes focusing on the y-axis from ∼0.01 to ∼100, suggesting that *λ* can remain invariant even when underlying metabolic and trophic parameters shift.

## 4. Discussion

This study contributes to growing evidence that metabolic scaling is not fixed, as proposed by the Metabolic Theory of Ecology (MTE) (1), but instead exhibits significant ecological plasticity in response to environmental drivers (9,11,12). We found that temperature and predation exert interactive effects on metabolic scaling, highlighting the importance of ecological context in shaping physiological traits. Specifically, temperature increased the metabolic scaling exponent in the absence of predators but reduced it in their presence— suggesting a strong interaction between thermal environment and predation pressure.

This result aligns with previous studies that demonstrated that predator cues can modulate the thermal sensitivity of metabolic scaling (9,12). In the presence of fish predators or their chemical cues, the metabolic rate of young, small individuals of the freshwater amphipod *Gammarus minus* increased more in response to increased temperature than did that of older, larger amphipods, thus decreasing the metabolic scaling exponent *b*. Size-selective predation apparently favors reduced temperature-related increases in activity and associated metabolic rates in more vulnerable larger amphipods. By contrast, in the absence of fish predators or their cues, the metabolic rate of older, larger amphipods increased more in response to increased temperature than did that of younger, smaller amphipods, thus increasing the metabolic scaling exponent *b* (9,11,12). This pattern likely reflects a trophic cascade triggered by predator removal, in which large adult amphipods, no longer suppressed by fish predation, occupy a size-refugium and exhibit higher densities and intraspecific resource competition that favor cannibalism. With fewer top-down constraints, more abundant adults redirect size-specific predation pressure onto smaller conspecifics. As a result, juveniles now face heightened mortality risk, not from fish, but from larger members of their own population—restructuring intra-guild predation causing the observed shifts in size structure and metabolic scaling.

A similar mechanism may be operating in our mesocosms. In the presence of fish predators, larger more conspicuous macroinvertebrates may have minimized their exposure by dampening an increase in their activity and associated metabolic rate in response to increasing temperature, whereas smaller less conspicuous macroinvertebrates were able to exhibit a stronger thermal response, thus decreasing the metabolic scaling exponent *b* (12). Macroinvertebrate communities in the tanks included individuals ranging from approximately 1 mm to 15 mm in body length, spanning multiple taxonomic groups (e.g., midges, beetles, damselflies, dragonflies). This wide range of body sizes emphasizes that both small, cryptic taxa and larger, conspicuous individuals were present and likely contributed to the observed size-dependent responses. However, in the absence of fish predators, the large, predatory macroinvertebrates may have had higher activity levels that posed a greater predation risk to small macroinvertebrates, leading to the small invertebrates having reduced activity levels and metabolic rates. Although the overall distribution of macroinvertebrate body sizes did not differ significantly among treatments, it is possible that large predatory invertebrates were more active or behaviorally dominant in fish-free environments (39).

These observations support a broader conceptual framework in which mortality risk— from either predators or conspecifics—can significantly influence metabolic scaling patterns (44). They also highlight the complexity of predicting physiological outcomes in multitrophic systems, where responses are shaped not only by abiotic conditions but also by the structure and behavior of interacting species. A mechanistic understanding of ecological scaling must therefore account for both environmental variation and indirect effects mediated through trophic interactions. At the community level, our results contribute to a limited but growing body of research exploring how size scaling patterns manifest across multiple interacting species. Yet this is critical for predicting how entire ecosystems will respond to climate change, as many core processes—such as energy flow and trophic efficiency—emerge from species interactions rather than strictly individual physiology.

In our experiment, we applied a community-level approach by examining the size– abundance slope (*λ*) as a proxy for energy distribution across body sizes. Unexpectedly, *λ* remained stable across treatments, despite clear treatment effects on metabolic scaling. This result is surprising because one might expect fish predation to reduce the number of large macroinvertebrates, thereby decreasing λ. Although, changes in individual metabolic rates— such as increased respiration under warming or the removal of larger individuals by predators—should cascade to influence ISD, we did not observe this.

Several hypotheses may help explain this apparent inconsistency. First, fish predation may not have been strongly size selective. Although sunfish (*Lepomis spp*.) are known to preferentially consume larger macroinvertebrate prey (45,46), prey selectivity is often size-dependent and can vary with the size or developmental stage of the fish. Smaller sunfish, limited by gape size, tend to consume smaller prey, while larger individuals are capable of targeting larger and a wider range of prey items (47). A lack of strong size-selectivity would also influence our interpretation of the temperature-related changes in metabolic scaling, which in other studies have been attributed in part to predator-induced reductions in growth or activity in larger individuals (9,11,12).

Second, an explanation for the consistent λ values across treatments may lie in the balance among key ecological parameters that shape ISDs such as *TTE*, *PPMR*, and the metabolic scaling *b*. While these factors may shift in response to environmental conditions like temperature and predation, their combined influence on energy flow could remain stable. This idea is supported by previous findings from marine systems, where neither *PPMR* nor *TTE* showed consistent variation across thermal gradients (48). Across a range of *b* values— from 0.3 to 1—combinations of *PPMR* and *TTE* often yield λ values within empirically observed ranges. In particular, plausible *TTE* values (around 0.05–0.15) align with ISD *λ* values between –1.5 and –2.5, regardless of *b*. This suggests that even if metabolic scaling varies with temperature or predator exposure, the broader structure of energy distribution across body sizes can remain unchanged. Such stability could result from compensatory adjustments. For instance, warming tends to reduce the biomass of larger consumers (49), which would tend to lower PPMR. At the same time, it can reduce TTE by decreasing the nutritional quality of prey (50). If the relative changes in *PPMR* and *TTE* are proportionate, their ratio—and therefore *λ*—can remain stable. However, unequal changes between the two can lead to steeper λ values by restricting energy transfer to higher trophic levels (51).

Additionally, predator feeding behavior may further influence *PPMR* under warming. If omnivorous predators like sunfish become more flexible in their foraging—shifting toward smaller or suboptimal prey—this can further lower realized PPMR, buffering *λ* against steeper declines even as top-down control weakens. Omnivory, in particular, has been shown to compress food chain length and reduce *PPMR* by broadening the trophic niche toward smaller prey sizes. What this suggests is that even when *b* varies under different environmental treatments, energy flow across the system may remain effectively unchanged. As a result, size distribution structure at the community level can remain stable, even as community-level metabolism shifts in response to temperature and predation pressure.

Third, the absence variation in λ across treatments points to the need for a deeper understanding of how temperature and predation shape individual size distributions (ISDs), particularly over short timescales. One consideration is that the 30-day exposure to predators may not have been long enough for top-down forces to produce detectable changes in community structure. While metabolic responses to temperature and predation risk can occur quickly (9), changes in population-level patterns—such as abundance distributions—often develop more gradually through processes like survival, growth, and reproduction (33,52). In this context, the stability in *λ* may reflect a delay in demographic responses rather than a true absence of ecological effect. While the temperature treatment may have underestimated the effect of warming on community structure, evidence from the experiment confirms that both temperature and predation treatments were ecologically active, as fish exhibited clear increases in body size, nearly doubling their mass (Table S1), and heated tanks consistently maintained elevated temperatures relative to ambient conditions (Figure S1).

Fourth, predator presence may have influenced species colonization or composition rather than across species size structure. Fish may deter or exclude certain taxa from colonizing or persisting in enclosures, not because of prey size per se, but due to species-specific behavioral or ecological traits. For example, fish-tolerant species may have broader size ranges or faster life histories that buffer the community against structural change. In this scenario, community composition could shift in response to predation without producing a change in λ, if the size distributions of new and excluded species were functionally redundant. Such species sorting processes have been observed in other aquatic systems, where predators alter community composition without systematically truncating size spectra (53,54).

Taken together, these hypotheses underscore the complexity of scaling relationships in ecological systems. While individual-level metabolic traits clearly respond to environmental variation, these responses do not necessarily translate to predictable changes in community structure. This finding adds to growing evidence that the MTE’s assumptions—particularly regarding the tight coupling of metabolism, body size, and abundance—are overly simplistic (37,55). This implies that community-level structure may be buffered against short-term physiological changes, perhaps due to compensatory shifts in species composition, colonization, or redundancy in size distribution (Figure 4). Such resilience highlights the need to link individual energetics with higher-order ecological structure to fully understand how ecosystems respond to environmental changes.

**Figure 4.**
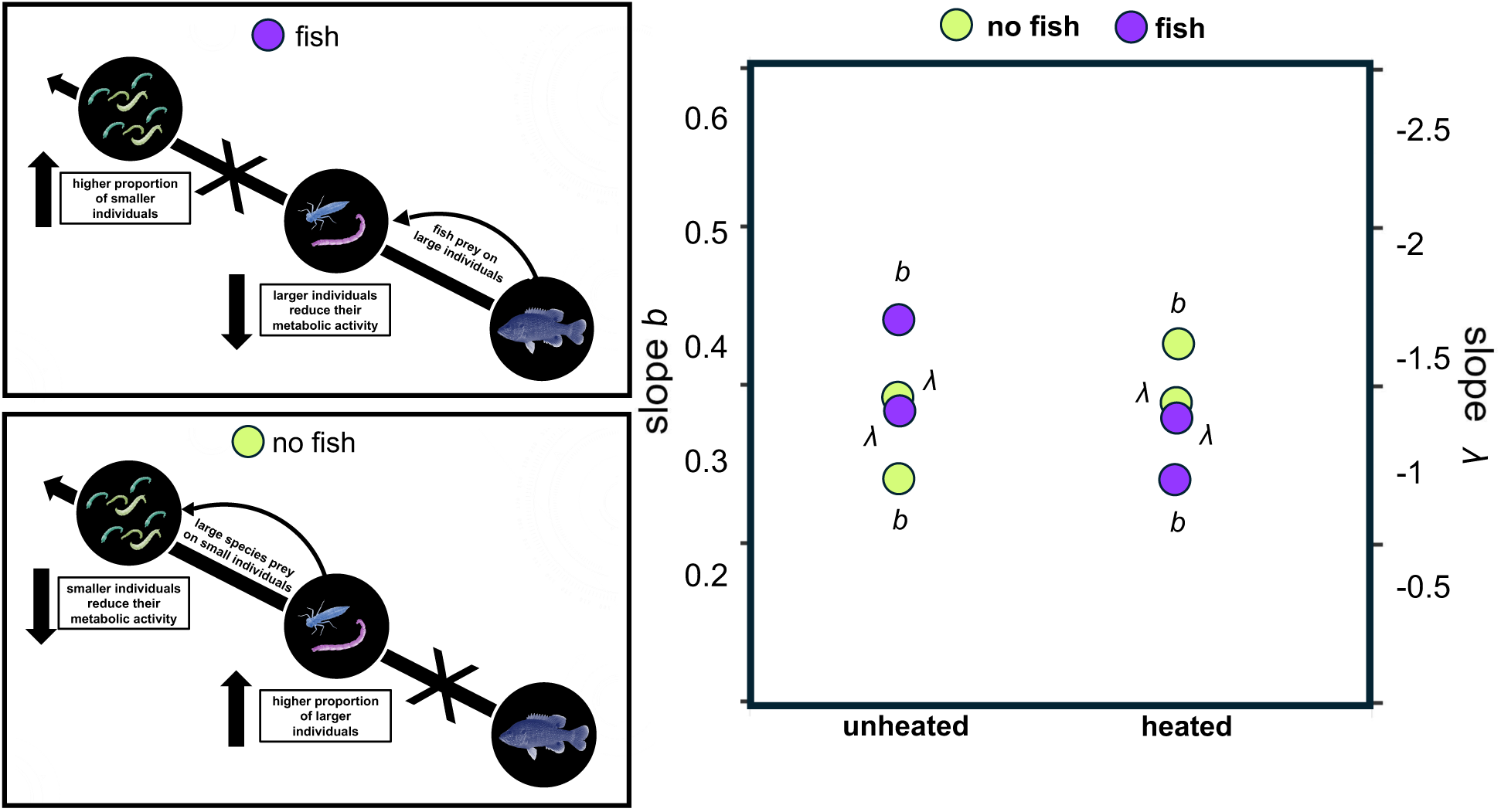
Conceptual diagrams (left) and empirical results (right) showing how fish predation and temperature influence size structure and metabolic scaling. In systems with fish (top left), predation on large individuals leads to a higher proportion of smaller individuals, while larger individuals reduce their metabolic activity. Without fish (bottom left), large invertebrates prey on smaller individuals, leading to a higher proportion of larger individuals, while smaller individuals reduce their metabolism. The scatter plot (right) shows slopes of metabolic scaling (*b*) and size distribution (*λ*) under heated and unheated conditions, with green and purple points representing fish and no-fish treatments, respectively. Results indicate that λ remains stable, while *b* varies with temperature and predation.

A critical insight from this study is that macroinvertebrates colonized the mesocosms naturally, resulting in a self-assembling, dynamic community more representative of real ecosystems than artificially assembled ones. This continuous recruitment, combined with natural community cycling and turnover, likely played a key role in buffering the system against structural disruptions from both warming and predation. In such open systems, ecological interactions—such as dispersal, recolonization, and species sorting—can maintain community composition and size structure even under substantial environmental stress (56–58). The resilience of the ISD’s *λ* observed across treatments may therefore reflect not only functional redundancy or trait compensation but also the stabilizing effect of ongoing immigration from regional species pools (59). These dynamics are rarely accounted for in closed experimental systems, where community responses can be more sensitive to perturbation. Our findings suggest that realistic colonization processes are crucial for understanding how ecosystems maintain structural integrity even as physiological responses vary at the individual level.

## 5. Conclusion

Overall, our findings highlight that *λ* is not solely determined by *b*. Instead, community-level structure appears to be maintained through trade-offs among key trophic parameters. Even when individual metabolic responses shift due to warming or predator presence, the broader energy distribution across body sizes may remain stable if underlying trophic efficiencies and interaction strengths adjust in tandem. Our findings reinforce the view that metabolic scaling is not a fixed, universal property, but one that is ecologically contingent—shaped by thermal conditions, species interactions, and behavioral responses to predation. The observed shifts in metabolic scaling under different predator regimes, coupled with the stability of community structure, suggest that physiological and structural responses can be decoupled, particularly in dynamic, self-assembling communities. This challenges core assumptions of the MTE and underscores the need for models that incorporate biotic context, mortality risk, and trophic interactions. Predicting ecological responses to climate warming will require moving beyond temperature in isolation and embracing the complexity of ecological networks and community assembly processes.

## Acknowledgments

This material is based upon work supported by the National Science Foundation under Grant Nos. 2106067 to JSW and 2106068 to JRJ. The National Ecological Observatory Network Biorepository at Arizona State University provided samples for organic matter collected as part of the NEON Program. The use of vertebrates was approved under the University of South Dakota Institutional Animal Care and Use Committee (#03-03-18-21). Computations supporting this project were performed on High Performance Computing systems at the University of South Dakota, funded by NSF Award OAC-1626516.

## Supplementary Information

**Table S1.** Growth of individual fish in each mesocosm.

| Treatment | Mesocosm | Dry Mass (g) |  |  |
| --- | --- | --- | --- | --- |
|  |  | Start | End | Change |
| +heat\|+fish | 1 | 1.8 | 5.2 | 3.4 |
|  | 4 | 5.1 | 5.2 | 0.1 |
|  | 9 | 2.2 | 4.6 | 2.4 |
|  | 15 | 2.1 | 4.5 | 2.3 |
|  | 18 | 2.3 | 5.3 | 3.0 |
|  | 19 | 1.9 | 3.5 | 1.6 |
| | Mean ± SD | 2.6 ± 1.1 | 4.7 ± 0.6 | 2.1 ± 1 |
| -heat\|+fish | 5 | 3.1 | 5.0 | 1.8 |
|  | 7 | 3.3 | 6.8 | 3.5 |
|  | 11 | 2.5 | 6.1 | 3.7 |
|  | 14 | 2.6 | 5.2 | 2.6 |
|  | 20 | 2.1 | 3.4 | 1.3 |
|  | 24 | 2.2 | 3.1 | 0.9 |
| | Mean ± SD | 2.6 ± 0.4 | 4.9 ± 1.3 | 2.3 ± 1 |

**Figure S1.**
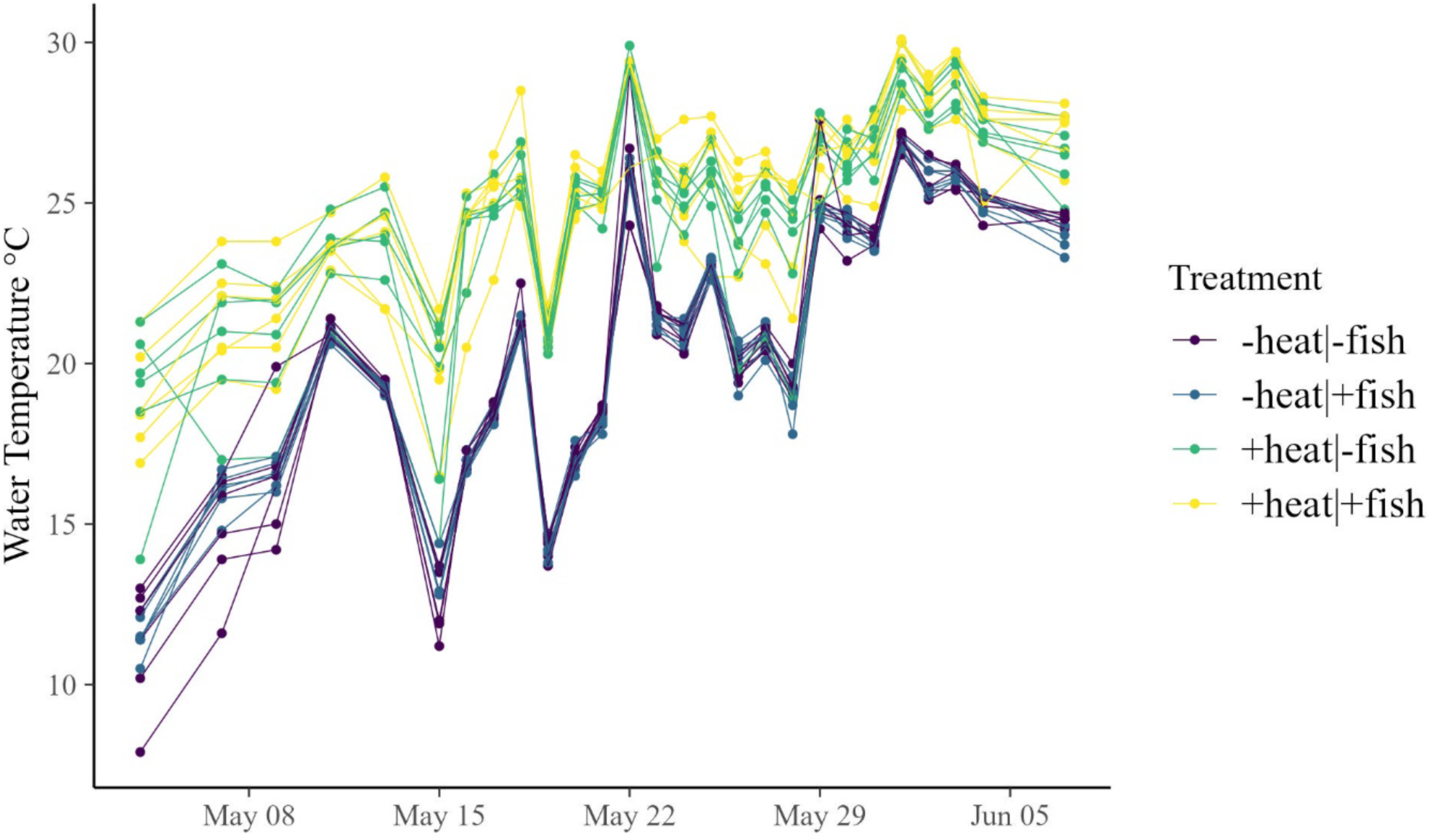
Temperature measurements over time in the experimental mesocosms (n = 24). Each dot represents a single temperature measurement from a mesocosm on a given day. Dates range from May 4 to June 7.

**Figure S2.**
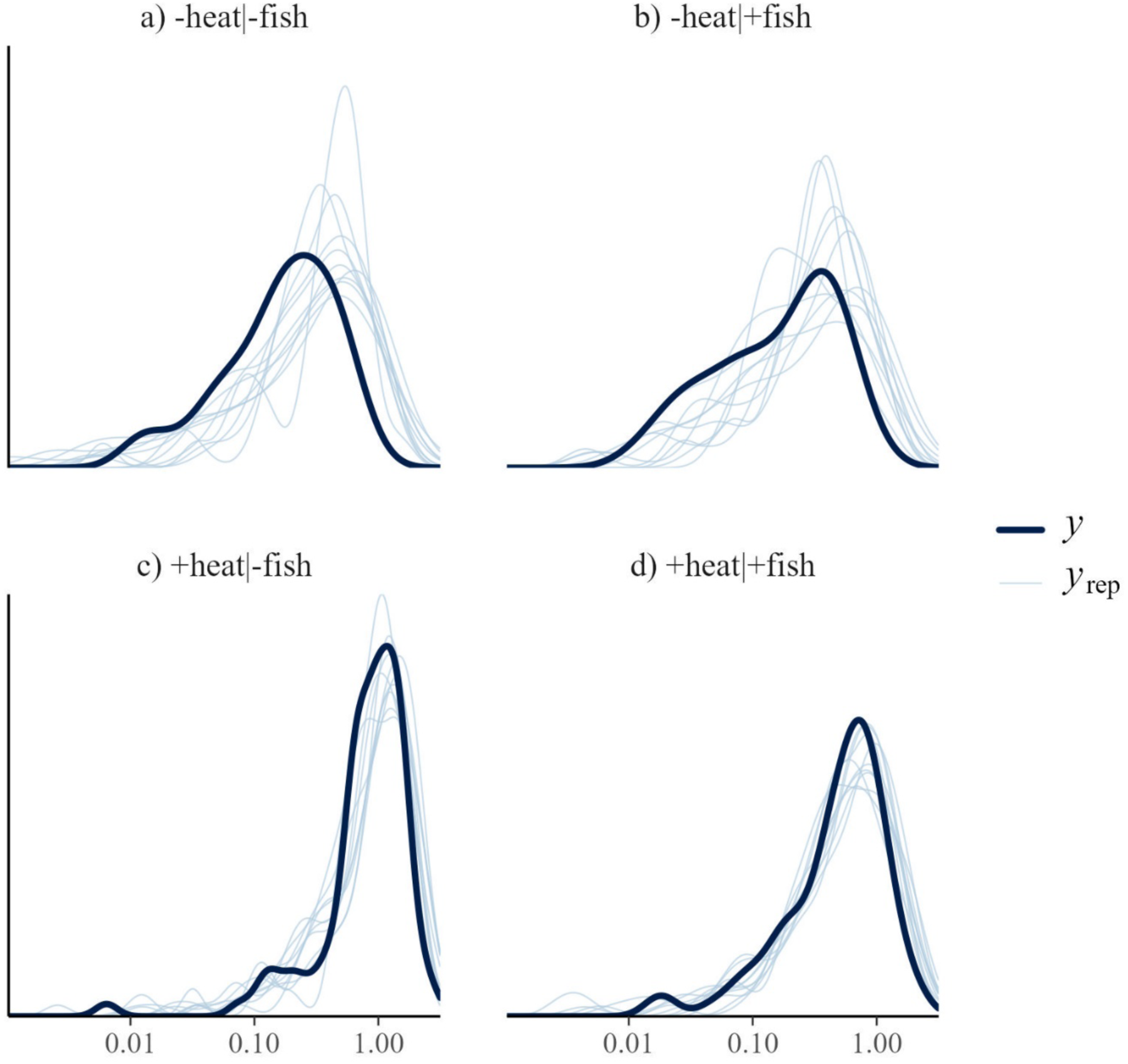
Model fit of metabolic scaling. For each of the four treatments, the density of raw data (*y*) is shown in dark blue. The lighter blue lines (*y*rep) show 10 datasets simulated from the posterior predictive distribution. The similarity in *y* and *y*rep indicate good model fit.

**Figure S3.**
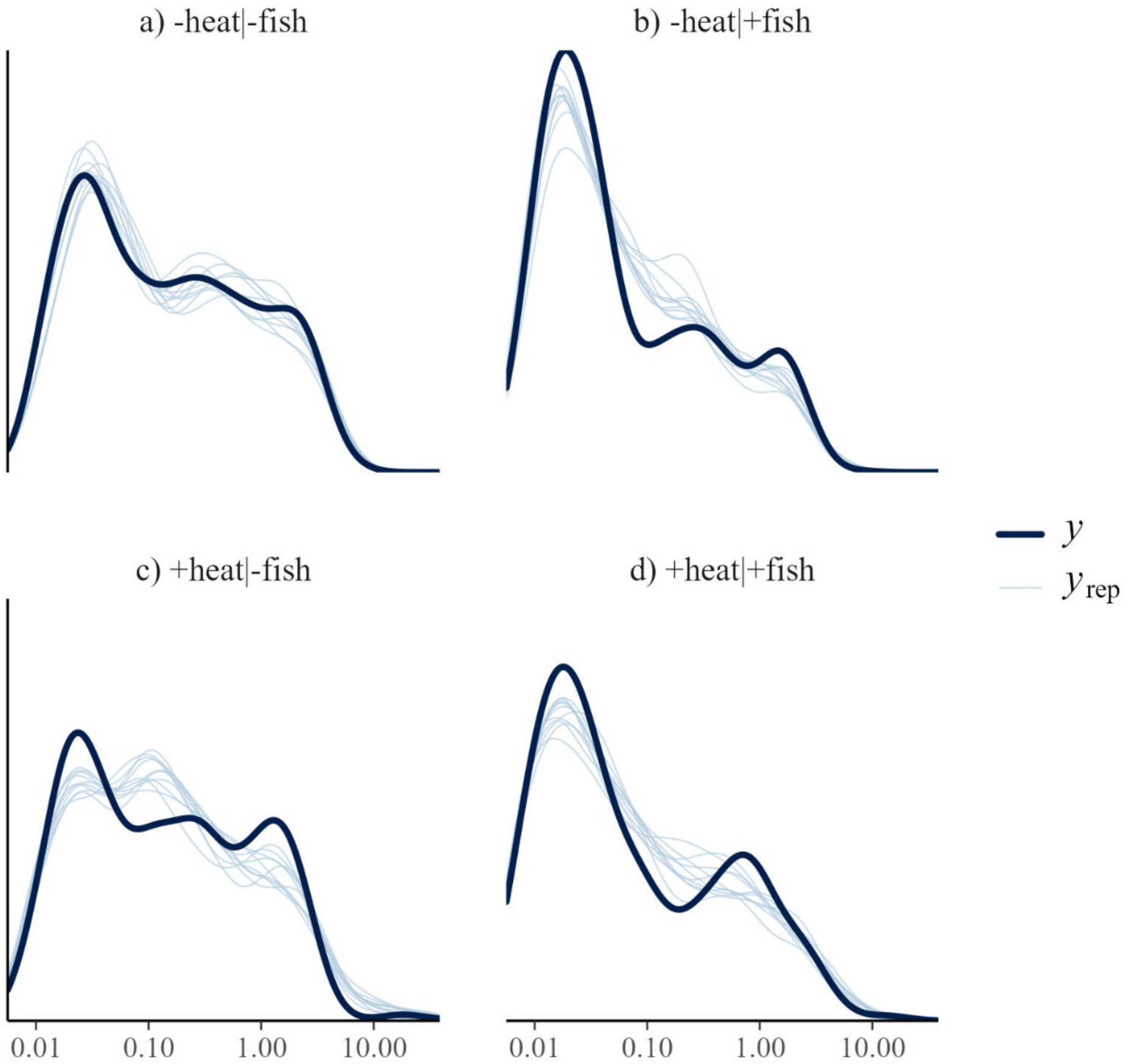
Model fit of the individual size distributions. For each of the four treatments, the density of raw data (*y*) is shown in dark blue. The lighter blue lines (*y*rep) show 10 datasets simulated from the posterior predictive distribution. The similarity in *y* and *y*rep indicate good model fit.

**Figure S4.**
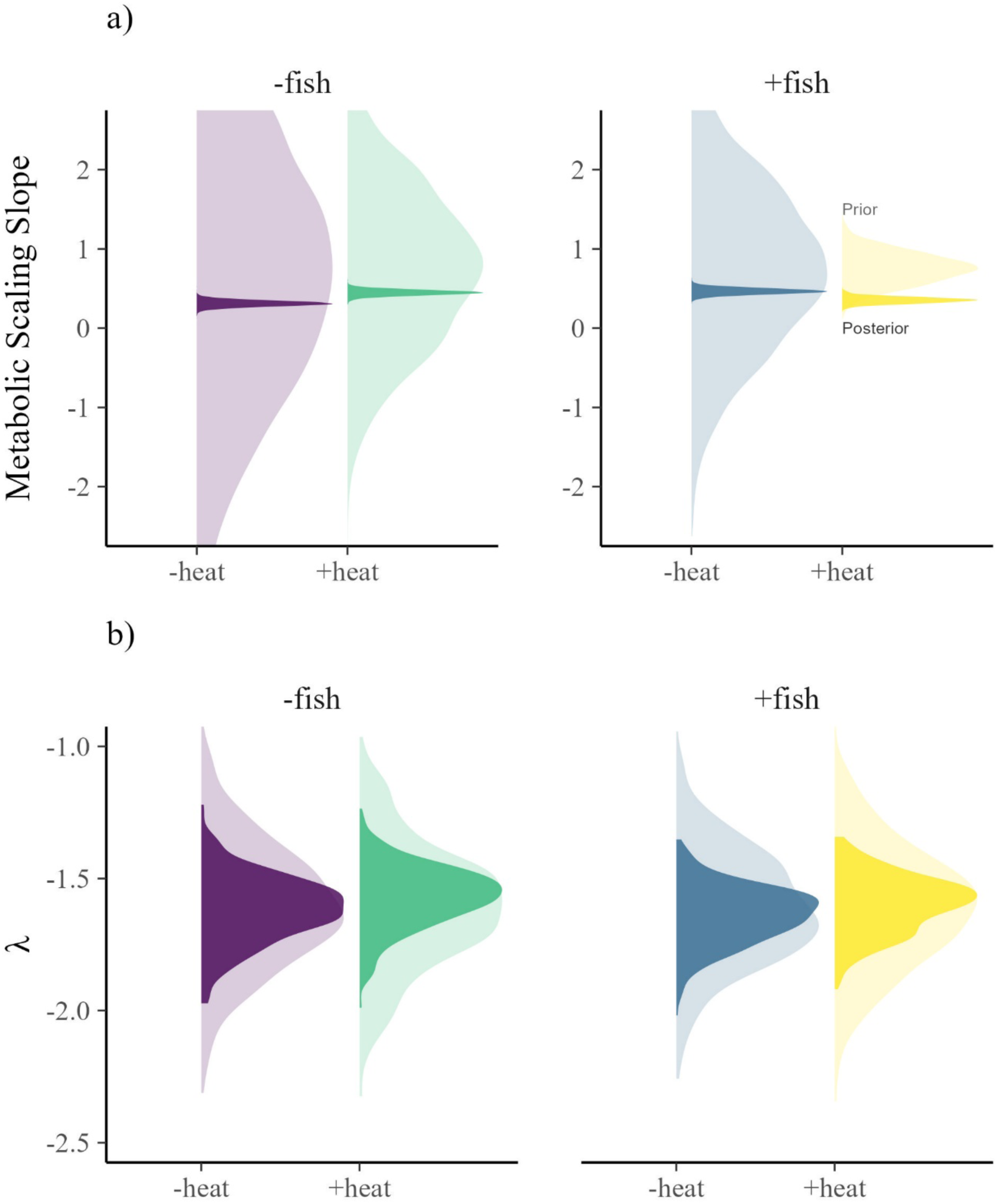
Comparison of the priors and posteriors. The dark colors show the posterior distributions (as in Figure 1). The lighter colors show the prior distributions. The difference in the prior and posterior is a proxy for the amount of information that the models learned from the data.

## References

1. Brown JH, Gillooly JF, Allen AP, Savage VM, West GB. 2004 Toward a metabolic theory of ecology. Ecology 85, 1771–1789.

2. Savage VM, Gillooly JF, Woodruff WH, West GB, Allen AP, Enquist BJ, Charnov EL. 2004 The predominance of quarter-power scaling in biology. Funct. Ecol. 18, 257– 282.

3. Gjoni V, Glazier DS, Wesner JS, Ibelings BW, Thomas MK. 2023 Temperature, resources and predation interact to shape phytoplankton size–abundance relationships at a continental scale. Glob. Ecol. Biogeogr. 32, 2006–2016.

4. Banavar JR, Moses ME, Brown JH, Damuth J, Rinaldo A, Sibly RM, Maritan A. 2010 A general basis for quarter-power scaling in animals. PNAS 107, 15816–15820.

5. Allen AP, Brown JH, Gillooly JF. 2002 Global biodiversity, biochemical kinetics, and the energetic-equivalence rule. Science 297, 1545–1548.

6. Glazier DS. 2005 Beyond the 3/4-power law: variation in the intra- and interspecific scaling of metabolic rate in animals. Biol. Rev. 80, 611–662.

7. White EP, Ernest SKM, Kerkhoff AJ, Enquist BJ. 2007 Relationships between body size and abundance in ecology. Trends Ecol. Evol. 22, 323–330.

8. Killen SS, Atkinson D, Glazier DS. 2010 The intraspecific scaling of metabolic rate with body mass in fishes depends on lifestyle and temperature. Ecol. Lett. 13, 184– 193.

9. Gjoni V, Basset A, Glazier DS. 2020 Temperature and predator cues interactively affect ontogenetic metabolic scaling of aquatic amphipods. Biol. Lett. 16, 20200267.

10. Glazier DS. 2022 Variable metabolic scaling breaks the law: from ‘Newtonian’ to ‘Darwinian’ approaches. Proc. R. Soc. B 289, 20221605.

11. Glazier DS, Butler EM, Lombardi SA, Deptola TJ, Reese AJ, Satterthwaite EV. 2011 Ecological effects on metabolic scaling: amphipod responses to fish predators in freshwater springs. Ecol. Monogr. 81, 599–618.

12. Glazier DS, Gring JP, Holsopple JR, Gjoni V. 2020 Temperature effects on metabolic scaling of a keystone freshwater crustacean depend on fish-predation regime. J. Exp. Biol. 223, jeb232322.

13. Damuth J. 1981 Population density and body size in mammals. Nature 290, 699–700.

14. Marquet PA, Navarrete SA, Castilla JC. 1990 Scaling population density to body size in rocky intertidal communities. Science 250, 1125–1127.

15. Enquist BJ, Brown JH, West GB. 1998 Allometric scaling of plant energetics and population density. Nature 395, 163–165.

16. Malerba ME, Marshall DJ. 2019 Size–abundance rules? Evolution changes scaling relationships between size, metabolism and demography. Ecol. Lett. 22, 1407–1416.

17. O’Gorman EJ, Zhao L, Pichler DE, Adams G, Friberg N, Rall BC, et al. 2017 Unexpected changes in community size structure in a natural warming experiment. Nat. Clim. Chang. 7, 659–663.

18. Dickie LM, Kerr SR, Boudreau PR. 1987 Size-dependent processes underlying regularities in ecosystem structure. Ecol. Monogr. 57, 233–250.

19. Jonsson T, Cohen JE, Carpenter SR. 2005 Food webs: from connectivity to energetics. Amsterdam, The Netherlands: Elsevier.

20. Reuman DC, Mulder C, Raffaelli D, Cohen JE. 2008 Three allometric relations of population density to body mass: theoretical integration and empirical tests in 149 food webs. Ecol. Lett. 11, 1216–1228.

21. Blanchard JL, Heneghan RF, Everett JD, Trebilco R, Richardson AJ. 2017 From bacteria to whales: using functional size spectra to model marine ecosystems. Trends Ecol. Evol. 32, 174–186.

22. Hatton IA, Heneghan RF, Bar-On YM, Galbraith ED. 2021 The global ocean size spectrum from bacteria to whales. Sci. Adv. 7, eabh3732.

23. Gillooly JF, Brown JH, West GB, Savage VM, Charnov EL. 2001 Effects of size and temperature on metabolic rate. Science 293, 2248–2251.

24. Schulte PM. 2015 The effects of temperature on aerobic metabolism: towards a mechanistic understanding of the responses of ectotherms to a changing environment. J. Exp. Biol. 218, 1856–1866.

25. Atkinson D. 1995 Effects of temperature on the size of aquatic ectotherms: exceptions to the general rule. J. Therm. Biol. 20, 61–74.

26. Forster J, Hirst AG, Atkinson D. 2012 Warming-induced reductions in body size are greater in aquatic than terrestrial species. PNAS 109, 19310–19314.

27. Saito VS, Perkins DM, Kratina P. 2021 A metabolic perspective of stochastic community assembly. Trends Ecol. Evol. 36, 280–283.

28. Daufresne M, Lengfellner K, Sommer U. 2009 Global warming benefits the small in aquatic ecosystems. PNAS, 12788–12793.

29. Yvon-Durocher G, Montoya JM, Trimmer M, Woodward G. 2011 Warming alters the size spectrum and shifts the distribution of biomass in freshwater ecosystems. Glob. Change Biol. 17, 1681–1694.

30. Pomeranz JPF, Junker JR, Wesner JS. 2022 Individual size distributions across North American streams vary with local temperature. Glob. Change Biol. 28, 848–858.

31. Arranz I, Grenouillet G, Cucherousset J. 2023 Human pressures modulate climate-warming-induced changes in size spectra of stream fish communities. *Nat*. Ecol. Evol. 7, 1072–1078.

32. Brose U et al. 2006 Consumer–resource body-size relationships in natural food webs. Ecology 87, 2411–2417.

33. Emmerson MC, Raffaelli D. 2004 Predator–prey body size, interaction strength and the stability of a real food web. J. Anim. Ecol. 73, 399–409.

34. Dell AI, Pawar S, Savage VM. 2014 Temperature dependence of trophic interactions are driven by asymmetry of species responses and foraging strategy. J. Anim. Ecol. 83, 70–84.

35. Shokri M, Marrocco V, Cozzoli F, Vignes F, Basset A. 2024 The relative importance of metabolic rate and body size to space use behavior in aquatic invertebrates. Ecol. Evol. 14, e11253.

36. Glazier DS. 2014 Metabolic scaling in complex living systems. Systems 2, 451–540.

37. Henry BL, Wesner JS. 2018 Severing ties: different responses of larval and adult aquatic insects to atrazine and selenium. Environ. Sci. Technol. 52, 8848–8857.

38. Wesner JS. 2010 Aquatic predation alters a terrestrial prey subsidy. Ecology 91, 1435– 1444.

39. Wesner JS, Pomeranz JPF, Junker JR, Gjoni V. 2024 Bayesian hierarchical modelling of size spectra. Methods Ecol. Evol. 15, 856–867.

40. Bürkner PC. 2017 brms: an R package for Bayesian multilevel models using Stan. J. Stat. Softw. 80, 1–28.

41. Gabry J, Simpson D, Vehtari A, Betancourt M, Gelman A. 2019 Visualization in Bayesian workflow. J. R. Stat. Soc. A 182, 389–402.

42. Wesner JS, Pomeranz JPF. 2021 Choosing priors in Bayesian ecological models by simulating from the prior predictive distribution. Ecosphere 12, e03739.

43. Glazier DS. 2025 Does death drive the scaling of life? Biol. Rev. 100, 586–619.

44. Mittelbach GG. 1981 Foraging efficiency and body size: a study of optimal diet and habitat use by bluegills. Ecology 62, 1370–1386.

45. Osenberg CW, Mittelbach GG. 1989 Effects of body size on the predator–prey interaction between pumpkinseed sunfish and gastropods. Ecol. Monogr. 59, 405–432.

46. Miller TJ, Crowder LB, Rice JA, Marschall EA. 1988 Larval size and recruitment mechanisms in fishes: toward a conceptual framework. Can. J. Fish. Aquat. Sci. 45, 1657–1670.

47. Barnes C, Maxwell D, Reuman DC, Jennings S. 2010 Global patterns in predator– prey size relationships reveal size dependency of trophic transfer efficiency. Ecology 91, 222–232.

48. Petchey OL, McPhearson PT, Casey TM, Morin PJ. 1999 Environmental warming alters food-web structure and ecosystem function. Nature 402, 69–72.

49. Cross WF, Hood JM, Benstead JP, Huryn AD, Nelson D. 2015 Interactions between temperature and nutrients across levels of ecological organization. Glob. Change Biol. 21, 1025–1040.

50. Jennings S, Mackinson S. 2003 Abundance–body mass relationships in size-structured food webs. Ecol. Lett. 6, 971–974.

51. Woodward G, Ebenman B, Emmerson M, Montoya JM, Olesen JM, Valido A, Warren PH. 2005 Body size in ecological networks. Trends Ecol. Evol. 20, 402–409.

52. White CR, Kearney MR. 2014 Metabolic scaling in animals: methods, empirical results, and theoretical explanations. Compr. Physiol. 4, 231–256.

53. Glazier DS. 2015 Is metabolic rate a universal ‘pacemaker’ for biological processes? Biol. Rev. 90, 377–407.

54. Harrison JF. 2017 Do performance–safety tradeoffs cause hypometric metabolic scaling in animals? Trends Ecol. Evol. 32, 653–664.

55. Leibold MA, Holyoak M, Mouquet N, Amarasekare P, Chase JM, Hoopes MF, et al. 2004 The metacommunity concept: a framework for multi-scale community ecology. Ecol. Lett. 7, 601–613.

56. Cadotte MW. 2006 Dispersal and species diversity: a meta-analysis. Am. Nat. 167, 913–924.

57. Holyoak M, Leibold MA, Holt RD. 2005 Metacommunities: Spatial Dynamics and Ecological Communities. Chicago, IL: University of Chicago Press.

